# Measuring the Contribution of Genomic Predictors to Improving Estimator Precision in Randomized trials

**DOI:** 10.1101/018168

**Authors:** Prasad Patil, Michael Rosenblum, Jeffrey T. Leek

## Abstract

The use of genomic data in the clinic has not been as widespread as was envisioned when sequencing and genomic analysis became common techniques. An underlying difficulty is the direct assessment of how much additional information genomic data are providing beyond standard clinical measurements. This is hard to quantify in the clinical setting where laboratory tests based on genomic signatures are fairly new and there are not sufficient data collected to determine how valuable these tests have been in practice. Here we focus on the potential precision gain from using the popular MammaPrint genomic signature in a covariate-adjusted, randomized clinical trial. We describe how adjustment of an estimator for the average treatment effect using baseline measurements can improve precision. This precision gain can be translated directly into sample size reduction and corresponding cost savings. We conduct a simulation study using genomic and clinical data gathered for breast cancer patients and find that adjusting for clinical factors alone provides a gain in precision of 5-6%, adjusting for genomic factors alone provides a similar gain (5%), and combining the two yields a 2-3% additional gain over only adjusting for clinical covariates.

## 1 Introduction

Incredible progress has been made in the characterization of the human genome, but the direct clinical benefit of these advances has not been clear for common complex diseases [5]. Problems with reproducibility [2], interpretability [12], and cost [1] have all slowed the translation of genomic markers to the clinic. One critical roadblock has been the uncertain value of genomic measurements for improving clinical practice [5]. A small number of laboratory tests based on genomic signatures have been approved for clinical use. Tests such as MammaPrint [20], Oncotype DX [14], and Prosigna [15] rely on measurement of expression for a set of genes that are associated with differential survival and severity of breast cancer cases.

It is difficult to evaluate the clinical value these genomic signatures add beyond standard clinical factors measured for all breast cancer patients, such as age, estrogen receptor status, tumor size, and tumor grade. It is also known that tests based on genomic signatures are not part of the standard of care in many cases [9, 5]. Ongoing clinical trials are being performed to evaluate the value of some of these signatures to make adaptive treatment decisions [3].

Here we propose to evaluate the use of genomic signatures in a different setting by considering the value added from using genomic signatures in designing a randomized clinical trial of a new treatment versus control. One of the principal goals of a randomized trial is to estimate the average treatment effect. We propose to use a genomic signature measured at baseline (pre-randomization) as a covariate that will be adjusted for in the analysis at the end of the randomized trial. If the genomic signature is prognostic for the primary outcome in the trial, this can lead to improved precision in estimating the average treatment effect [8].

Our aim is to determine the additional prognostic value of the genomic signature beyond the outcome variability already explained by standard clinical baseline variables. We describe a measure of prognostic value that directly translates into a reduction in the sample size required to achieve a desired power in a randomized trial. Prior work exists on using baseline variables to improve precision in the analysis of randomized trials (called covariate adjustment) [21, 22, 7, 19, 17, 10]. To the best of our knowledge, the value added by a genomic signature in this context has not yet been assessed.

We present results from a Monte Carlo study based on the data used to validate the MammaPrint model [6]. We compare the precision gain when clinical covariates are supplemented with genomic predictions to the gain when using clinical covariates alone to assess how much additional value is provided by the genomic data. This technique allows us to approximate the added value of certain genomic predictions in increasing precision in the analysis of randomized trial data.

## 2 Methods

### 2.1 Data

Microarray data used to validate the MammaPrint model [6] were gathered as described by the appendix of [13]. This dataset consists of 307 breast cancer patients and is characterized in Table 1. This table describes the key clinical factors gathered for these patients as well as their MammaPrint risk prediction, which is a classification based on the risk score calculated by the MammaPrint model [20]. We dropped 11 patients whose estrogen receptor (ER) status or tumor grade were unknown and conducted our analysis using the 296 remaining patients.

**Table 1:**
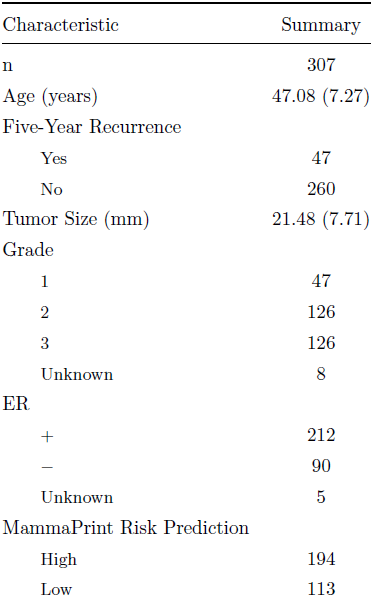
Baseline characteristics of curated dataset. Abbreviations: ER - estrogen receptor status, Grade - tumor severity grading (3 is most severe), Five-Year Recurrence - whether or not cancer has reappeared after five years, MammaPrint risk prediction - high or low risk for cancer recurrence. Age, Tumor Size are given as means with standard deviations.

| Characteristic | Summary |
| --- | --- |
| n | 307 |
| Age (years) | 47.08 (7.27) |
| Five-Year Recurrence |  |
| Yes | 47 |
| No | 260 |
| Tumor Size (mm) | 21.48 (7.71) |
| Grade |  |
| 1 | 47 |
| 2 | 126 |
| 3 | 126 |
| Unknown | 8 |
| ER |  |
| + | 212 |
| − | 90 |
| Unknown | 5 |
| MammaPrint Risk Prediction |  |
| High | 194 |
| Low | 113 |

We also provide three external breast cancer data sets to supplement our main result derived from the MammaPrint validation data. These datasets are described in the **Supplement**.

### 2.2 Statistical Method to Adjust for Baseline Covariates

We define the average treatment effect to be the difference between the population mean of the primary outcome under assignment to treatment and the population mean under assignment to control. The term “covariate adjustment” means that information from baseline variables is used to improve the precision in estimating the average treatment effect. This is done by adjusting for chance imbalances in baseline variables between treatment and control arms. Since our focus is improved precision for estimating the average treatment effect, we do not consider effects within subgroups; investigating the latter is an area for future research.

Increased precision for estimation of the average treatment effect can lead to trials with fewer participants and shorter duration if analysis timing is based on information monitoring [11, Chapter 7]. This means that analyses take place when preplanned amounts of information have accrued, where information is defined as the reciprocal of the estimator variance. Our definition of precision gain in Section 2.4 equals the percent sample size reduction from using an adjusted estimator compared to the unadjusted estimator (defined below), when information monitoring is used. This can equivalently be thought of as the percent sample size reduction to achieve a desired power at a local alternative, comparing these two estimators, asymptotically. These gains occur even when the treatment effect is zero, which is the setting of our simulation study.

Each participant in the trial contributes a data vector *D* = (*W, A, Y*), where *W* = (*W*_1_, . . . , *W*_*j*_) is a vector of covariates measured at baseline, *A* is an indicator of study arm (0 = control, 1 = treatment), and *Y* is a binary outcome of interest which in our case is the indicator of cancer recurrence within 5 years from baseline. The trial data consist of *n* independent, identically distributed participant data vectors 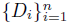 drawn from unknown distribution *P*. We assume a nonparametric model except that *W* and *A* are independent by randomization, and we assume the regularity conditions in [18, Section 2.2].

The goal is to estimate the average treatment effect,

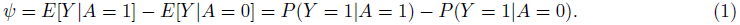

The unadjusted estimator of *ψ* is defined as

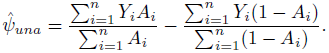

This estimator is consistent (i.e., converges in probability to the population average treatment effect *ψ*) but ignores the baseline variables *W*. If *W* is prognostic for *Y* then it is possible to improve precision by appropriately adjusting for *W*. Throughout, we do not assume that *W* contains information about treatment effect heterogeneity, i.e., who benefits more or less from treatment; we only use *W* as prognostic variables that explain some of the variation in *Y*. This variation could be unrelated to treatment.

To leverage the information in *W*, we apply two approaches: an enhanced efficiency, doubly-robust estimator of Rotnitzky *et. al.* [18], and a special case of their class of estimators that is slightly modified for use in the randomized trial context by Colantuoni and Rosenblum [8, Section 4.2]. We call these 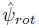 and 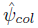, respectively. Software to compute the former estimator using R is given by [18], and software to compute the latter estimator is given in R and SAS by [8].

These estimators use parametric working models for the mean of the outcome given baseline variables and study arm. We call these working models since we do not assume they are correctly specified. The true data generating distribution may differ arbitrarily from the functional form of the model.

Computation of 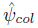 is accomplished via the following steps:

1. Let *α* = (*α*_0_, . . . , *α*_*j*_)*^T^*. Fit the following propensity score working model for *P*(*A* = 1*|W*): *g*(*W, α*) = logit ^*−1*^(*α*_0_+ *α*_1_*W*_1_+ . . .+ *α*_j_*W*_j_) via maximum likelihood parameter estimation. From this model fit, obtain estimates 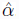.
2. For each arm *a* ∈ {0,1}, define the working model 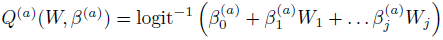 for *E*(*Y |A* = *a, W*). Fit the above model for *a* = 1 using weighted logistic regression with weights 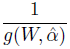 and using only participants with *A* = 1 to obtain estimated coefficients 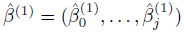. Define the initial estimator for 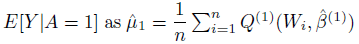, where the sum is taken over all participants. The estimator 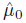 for *E*[*Y |A*= 0] is obtained analogously by setting *a* = 0, replacing *A* = 1 with *A* = 0, and replacing 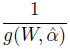 by 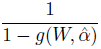 above.
3. Define the new covariate 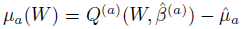 for each a ∈ {0, 1}, which uses 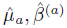 as estimated in step 2. Fit the following augmented propensity score model for *P*(*A* = 1|*W*): *g*_aug_(*W, α, γ*) = logit^−1^(*α*_0_+ *α*_1_*W*_1_+ . . . + *α_j_W_j_*+ *γ*_0_*µ*_0_(*W*)+ *γ*_1_*µ*_1_(*W*)) using maximum likelihood estimation to obtain estimated coefficients 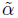 and 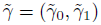.
4. Recompute step 2 using 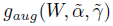 in place of 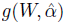 in the weights to obtain new estimates 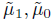. Define the average treatment effect estimator 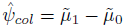

The estimator 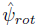 of [18, Section 2] is as above except that it involves solving a non-convex optimization problem leading to additional covariates in the model *g*_*aug*_ in step 3, which leads to enhanced efficiency guarantees. A link to the R code implementing both estimators is provided in Section 2.5.

Throughout, we assume there is no missing data and we observe the vector (*W*_*i*_, *A*_*i*_, *Y*_*i*_) for each participant *i*. We also assume outcomes are observed soon after enrollment. By the randomization assumption, the models *g* and *g*_*aug*_ are correctly specified as long as each contains an intercept; however, the models *Q*^(0)^, *Q*^(1)^ will typically be misspecified. An important feature of the above estimators is that they are consistent regardless of whether the parametric models *Q*^(0)^, *Q*^(1)^ are correctly specified; that is, consistency holds even when the true data generating distribution *E*(*Y* |*A* = *a, W*) does not have the form *Q*^(*a*)^(*W, β*^(*a*)^) for any *β*. Furthermore, each of these adjusted estimators is guaranteed to have asymptotic precision equal to or greater than that of the unadjusted estimator.

It is also possible to use the output of step 2 to directly construct the estimator 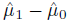 of the average treatment effect. This estimator is called the double-robust weighted least squares estimator (DR-WLS) and is attributed to Marshall Joffe by Robins *et al.*[16]. The value of adding steps 3 and 4 is that the resulting estimator has been proved to be asymptotically as or more precise than the unadjusted estimator [18, 8].

### 2.3 Prognostic Covariates

The baseline variables *W* used in the estimators defined above must be pre-specified. They can be any functions of measurements made prior to randomization. We define four sets of covariates of interest which we will adjust for using the procedure described in Section 2.2:

- *W*_−*ER*_: {Age, Tumor Size, I(Tumor Grade = 2), I(Tumor Grade = 3)}
- *W*_*C*_: {Age, Tumor Size, I(Tumor Grade = 2), I(Tumor Grade = 3), ER Status}
- *W*_*G*_: {MammaPrint Risk Category}
- *W*_*CG*_: {Age, Tumor Size, I(Tumor Grade = 2), I(Tumor Grade = 3), ER Status} MammaPrint Risk Category}

Here, *I*(*TumorGrade* = 2) is an indicator of whether or not the patient’s tumor is severity grade 2.

With these four sets of covariates, we are able to contrast gains in precision from different covariate sources. We may compare adjusting for *W*_−*ER*_ and *W*_*C*_ to determine how much adding a clinical covariate (ER status) to other clinical covariates improves precision. We can also determine the raw value of the genomic predictor of interest (*W*_*G*_) as well as the comparative gain from the genomic predictor over clinical covariates (*W*_*CG*_).

We have chosen to consider the clinical covariates stated here because they reflect quantities that clinicians commonly use to evaluate cancer-related risks and courses of therapy. The number of covariates we are adjusting for here exceeds what is recommended in [8]- they recommend 2-3 adjustment covariates. This means that we risk non-negligible increases in estimator variance if our covariates do not provide information about the outcome. We weigh this risk with the desire to compare the value of MammaPrint to the full set of relevant clinical covariates available in our dataset, and present an examination of the potential losses in the Results section.

### 2.4 Monte Carlo Trial Simulation

We conducted a simulation study with the goal of comparing the variance of the unadjusted and adjusted estimators to determine how much precision we may gain from adjusting for clinical and genomic covariates. We constructed data generating distributions that mimic features from our real data set.

To preserve the relationship between outcome and potentially prognostic covariates from the original data set, we resample participants with replacement and create a new 296-patient sample for each simulated trial; we record (*W, Y*) for each resampled participant. This maintains the empirical joint distribution of (*W, Y*), preserving the correlation of these variables. In each simulated trial, the study arm assignment *A* of each participant is a random draw from Bernoulli distribution with probability 1*/*2 of being 0 or 1, independent of (*W, Y*). The population average treatment effect defined in (1) corresponding to the above data generating distribution is therefore *ψ* = 0.

Using resampling as described above, we construct *I*= 10,000 simulated trial datasets (each of sample size 296). Using the *i*^*th*^ simulated dataset, we compute the unadjusted treatment effect estimator 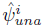 and compute the adjusted estimators 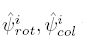 using each of the four predefined covariate sets *W*_*−ER*_, *W*_*C*_, *W*_*G*_, *W*_*CG*_. We then approximate the bias and variance of each of these estimators using their observed distribution over the 10,0000 replicated trials. Since *ψ* = 0, the bias *B* of an estimator is its average value over infinitely many hypothetical trials; we approximate this by the average over the 10,000 simulated trials we conducted. We similarly approximate the variance. For the unadjusted estimator, 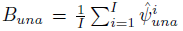, 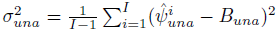. The bias and variance approximations for the Rotnitzky and Colantuoni estimators are calculated similarly. We are able to estimate the bias directly in the form *B*_*una*_

We define the percent precision gains (called precision gains below) due to the use of each adjusted estimator in comparison to the unadjusted estimator, as approximated by simulation, as 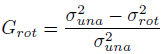 x 100 and 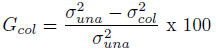 The percent precision gain equals, asymptotically (as sample size goes to infinity), the percent reduction in sample size to achieve a desired power comparing the adjusted versus unadjusted estimators. It equals 1 *−*1*/RE*, where *RE* is the asymptotic relative efficiency.

Simulations were conducted via the `BatchJobs` R package [4], which allows for an interface between R and a cluster queuing system. We parallelized such that 100 simulated datasets were constructed concurrently by each of 100 processors on a Sun Grid Engine (SGE) cluster, which sped up the computation of our approximations accordingly.

We conducted an additional set of simulations to examine how sensitive the above simulation study is to the particularities of the original dataset. In this second setting, we take ten random subsamples of size *m* = 222 (75% of our sample size) of the original dataset. For each of these subsamples, we resample with replacement a dataset of size 296 - our original sample size - and do this resampling 10,000 times. We then calculate *G*_*rot*_, *G*_*ros*_ for the different adjustment covariate scenarios exactly as described above for each subsample. This gives us ten realizations of *G*_*rot*_, *G*_*ros*_. We present a histogram of these realizations for our cases of interest (clinical, clinical + genomic) to show how much variation is caused by slightly changing the original data set.

### 2.5 Reproducibility

All analyses presented here are completely reproducible. Code and data files are available at https://github.com/leekgroup/genesigprecision

## 3 Results

Table 2 presents the main results of our simulation procedure. We are able to approximate the percent gain in precision due to covariate adjustment when clinical and genomic predictions are incorporated either separately or together. Additional adjustment for genomic predictions via the MammaPrint Risk Score for cancer recurrence led to improved precision beyond what was achieved by adjusting for clinical covariates alone.

**Table 2:** Precision gain under different covariate adjustments. This table presents the simulated estimates for the treatment effect and variance of the treatment effect estimator when unadjusted 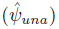 and under the two adjustment approaches 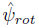, 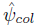. Each of 10,000 times, we resampled records from the original dataset with replacement to generate a new dataset of size *n* = 296. In every iteration, we adjusted the treatment effect estimator using a prespecified set of baseline covariates: *W*_−*ER*_ is clinical covariates only, excluding ER status; *W*_*C*_ is all clinical covariates only; *W*_*G*_ is only genomic covariates; *W*_*CG*_ includes all clinical and genomic covariates.

| Covariates | $B_{una}$ | $\sigma_{una}^2$ | $B_{rot}$ | $\sigma_{rot}^2$ | $B_{col}$ | $\sigma_{col}^2$ | $G_{rot}$ | $G_{col}$ |
| --- | --- | --- | --- | --- | --- | --- | --- | --- |
| $W_{-ER}$ | -0.00079 | 0.00182 | -0.00084 | 0.00172 | -0.00094 | 0.00172 | 5.54% | 5.14% |
| $W_C$ | -0.00079 | 0.00182 | -0.00101 | 0.00173 | -0.00087 | 0.00171 | 4.86% | 5.58% |
| $W_G$ | -0.00079 | 0.00182 | -0.00088 | 0.00172 | -0.00088 | 0.00172 | 5.36% | 5.36% |
| $W_{CG}$ | -0.00079 | 0.00182 | -0.00097 | 0.00167 | -0.00093 | 0.00169 | 7.78% | 6.96% |

The improvement due to inclusion of ER status above other clinical covariates was variable depending on adjustment method - *G*_*rot*_ was slightly negative, relatively, when ER status was included, while *G*_*col*_ showed a modest gain. The additional gain due to the genomic predictor exceeded this threshold. We found that the MammaPrint Score provided an additional precision gain of about 2-3% above using all clinical factors. *G*_*rot*_, *G*_*col*_ are also our approximations of by how much we would be able to reduce the sample size of our trial if we adjusted for the different sets of covariates when estimating the treatment effect.

To examine the impact of having a slightly different dataset than the original dataset, we created 10 modified datasets, each consisting of a random subset of 75% of the original participants (where each participant’s data vector was kept intact). Then, for each modified dataset, we ran the entire analysis described above, involving resampling 10,000 simulated trials each with 296 participants, as described at the end of section 2.4. Histograms of the gains due to clinical factors and the gains due to adding genomics appear in Figure 1. We found that the approximation of the gain may vary *±*3% around its mean, suggesting that our approximations from the resampling of the original datasets are fairly stable.

**Figure 1.**
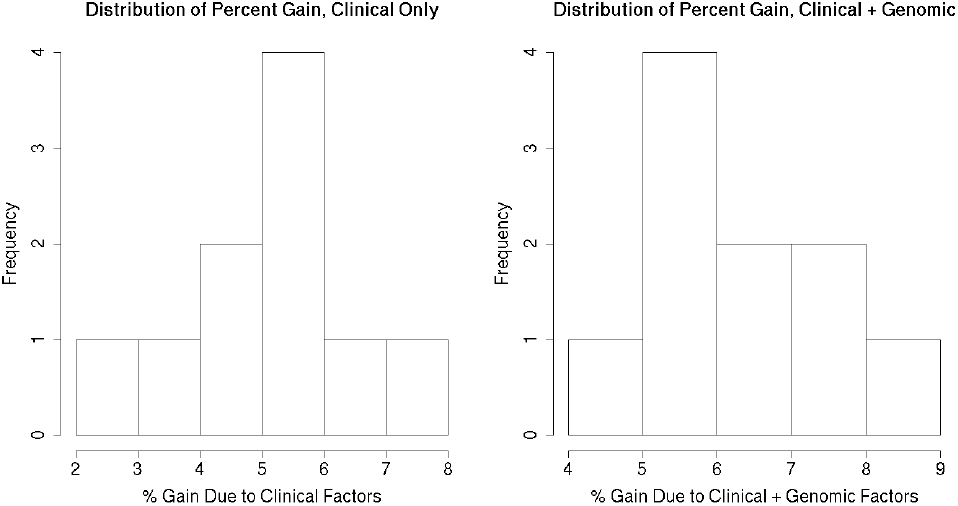
Variability in Percentage Gain. To approximate the variability in the percentage gain due to covariate adjustment, we took ten subsamples of 75% of our dataset and re-ran 10,000 resamplings on each of the ten. The left panel shows the histogram of the gain due to clinical covariates only (*W*_*C*_), and the right panel shows the gain due to both clinical and genomic covariates together (*W*_*CG*_) for the ten subsamples. We see that the gain may vary *±*3% from the average approximation we saw over the entire *I* = 10, 000 resampled datasets from the original run described in **Table 2**.

As an additional form of validation, we re-ran the same 10,000 resampling procedure described at the beginning of section 2.4 using three external datasets. The datasets and full results appear in the **Supplement**. These results showed either slight losses or gains comparable to those that we saw in the MammaPrint dataset when we included the MammaPrint risk prediction as a covariate.

Finally, we set a baseline for what we may expect in terms of precision gain when covariates provide no predictive value by completely permuting the labels in our dataset. This permutation should remove association between the outcome and covariates of interest, and we should see some loss due to adjusting for uninformative covariates. The results are shown in Table 3. As expected, all combinations of covariates produce losses near or below zero when labels are permuted and everything is independent. We observe a maximum loss in precision of about 4.6%, and this is certainly non-negligible, especially given that our gains in Table 2 are on the same order of magnitude. We stress that this is due to the inclusion of greater than the recommended number of adjustment covariates, as described in section 2.3. The potential losses are smaller if fewer uninformative covariates are used, or if the sample size is larger. We exceeded the recommended number of covariates to make a fair comparison between common clinical covariates and the addition of genomic predictions.

**Table 3:** Precision gain under different covariate adjustments - full permutation. This table presents the simulated estimates for the treatment effect and variance of the treatment effect estimator when unadjusted 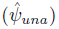 and under the two adjustment approaches 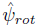,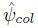. Each of 10,000 times, all covariate values in the patient data were permuted to simulate independence and a new treatment indicator was randomly generated. This was done to demonstrate that covariates independent of the outcome will provide no precision gain yet will not materially hurt precision. In every iteration, we adjusted the treatment effect estimator using a prespecified set of baseline covariates: *W*_−*ER*_ is clinical covariates only, excluding ER status; *W*_*C*_ is all clinical covariates only; *W*_*G*_ is only genomic covariates; *W*_*CG*_ includes all clinical and genomic covariates.

| Covariates | $B_{una}$ | $\sigma_{una}^2$ | $B_{rot}$ | $\sigma_{rot}^2$ | $B_{col}$ | $\sigma_{col}^2$ | $G_{rot}$ | $G_{col}$ |
| --- | --- | --- | --- | --- | --- | --- | --- | --- |
| $W_{-ER}$ | -0.00088 | 0.00354 | -0.00045 | 0.00368 | -0.00073 | 0.00363 | -4.02% | -2.58% |
| $W_C$ | -0.00088 | 0.00354 | -0.00074 | 0.00370 | -0.00056 | 0.00368 | -4.56% | -3.81% |
| $W_G$ | -0.00088 | 0.00354 | -0.00091 | 0.00356 | -0.00091 | 0.00356 | -0.4% | -0.4% |
| $W_{CG}$ | -0.00088 | 0.00354 | -0.00056 | 0.00372 | -0.00067 | 0.00369 | -4.9% | -4.2% |

## 4 Conclusion

Adjusting for baseline covariates and estimating treatment effects with augmented outcome regression models is one way to potentially improve treatment effect estimation precision. If baseline factors are collected for patients enrolled in a study, then adjusting for them can reduce the sample size necessary to obtain a desired precision in estimation of the average treatment effect. We showed via simulation that adjusting for the clinical covariates Age, Tumor Size, Tumor Grade, and ER status led to a gain of 5-6% in estimator precision. This percent gain can be directly translated into the amount (5%) by which sample size can be reduced when the adjusted estimator is used as compared to the unadjusted estimator.

Adjusting for genomic covariates on their own provides a similar precision gain of 5-6%. The precision gain when adjusting for genomic covariates in addition to standard clinical variables was 1-2%, resulting in similar reductions in sample size. This gain is modest, but not surprising given the prognostic power of the MammaPrint score in the validation set examined here (89% sensitive to high risk-of-recurrence patients, 42% specific to low risk-of-recurrence [13]). This gain is comparable to the gain one would get by including ER status as a baseline adjustment variable in addition to the other clinical covariates. A cost/benefit analysis taking into account the cost of running the MammaPrint test and the relative gain it may impart is warranted. The approach described here allows one to quantify the value of using MammaPrint along with clinical variables for improving estimator precision in a clinical trial.

